# Association between plasma phospho-tau181 and cognitive change from age 73 to 82: Lothian Birth Cohort 1936

**DOI:** 10.1101/2021.11.24.469836

**Authors:** Tyler S. Saunders, Amanda Heslegrave, Declan King, Sarah Harris, Craig Ritchie, Graciela Muniz-Terrera, Ian J. Deary, Simon R. Cox, Henrik Zetterberg, Tara Spires-Jones

**Affiliations:** UK Dementia Research Institute and Centre for Discovery Brain Sciences at the University of Edinburgh, Edinburgh, EH8 9JZ, United Kingdom; UK Dementia Research Institute at University College London, UCL Institute of Neurology, Queen Square, London, WC1N 3BG, United Kingdom; Department of Neurodegenerative Disease, UCL Institute of Neurology, Queen Square, London, WC1N 3BG, United Kingdom; Lothian Birth Cohort studies, Department of Psychology, University of Edinburgh, Edinburgh, EH8 9AD, United Kingdom; Edinburgh Dementia Prevention & Centre for Clinical Brain Sciences, University of Edinburgh, Edinburgh, EH4 2XU, United Kingdom; Department of Psychiatry and Neurochemistry, Sahlgrenska Academy at the University of Gothenburg, S-431 80 Mölndal, Sweden; Clinical Neurochemistry Laboratory, Sahlgrenska University Hospital, S-431 80 Mölndal, Sweden; Hong Kong Center for Neurodegenerative Diseases, Clear Water Bay, Hong Kong, China

**Keywords:** Tau proteins, Biomarkers, Cognition, Blotting, Western, Plasma P-tau181, Array Tomography

## Abstract

**INTRODUCTION:** Plasma phospho-tau 181 (p-tau181) is a promising blood biomarker for Alzheimer’s disease. However, its predictive validity for age-related cognitive decline without dementia remains unclear. Several forms of p-tau have been shown to contribute to synapse degeneration, but it is unknown whether p-tau181 is present in synapses. Here, we tested whether plasma p-tau181predicts cognitive decline and whether it is present in synapses in human brain.

**METHODS:** General cognitive ability and plasma p-tau181 concentration were measured in 195 participants at ages 72 and 82. Levels of p-tau181 in total homogenate and synaptic fractions were compared with western blot (n=10-12 per group), and synaptic localisation was examined using array tomography.

**RESULTS:** Elevated baseline plasma p-tau181 and increasing p-tau181 over time predicted steeper general cognitive decline. We observe p-tau181 in neurites, presynapses, and post-synapses in the brain.

**DISCUSSION:** Baseline and subsequent change in plasma p-tau181 may represent rare biomarkers of differences in cognitive ageing across the 8^th^ decade of life and may play a role in synaptic function in the brain.

## 1. Introduction

By the year 2050, the world’s population of people aged over 60 is expected to reach 2 billion [1]. With an ageing population comes challenges, including that of age-related cognitive decline which can seriously limit an individual’s independence and their quality of life [2,3]. Biomarkers associated with age-related cognitive decline will assist in understanding its underlying mechanisms [4] and may eventually improve prognoses.

Hyperphosphorylated tau (p-tau) accumulation and spread in the brain is a pathological hallmark of neurodegenerative diseases, such as Alzheimer’s disease (AD) and other tauopathies [5,6]. Tau phosphorylated at site threonine^181^ (p-tau181) has been investigated as a biomarker in the cerebrospinal fluid (CSF) and blood in studies of AD [7–11]. Both CSF and plasma p-tau181 concentrations are elevated in AD relative to controls [12] and have been found to correlate with global cognitive function in those with mild cognitive impairment (MCI) [13].

Despite tau accumulation being present in typical ageing [14,15] and being associated with pre-morbid cognitive function [16], few studies have examined the utility of plasma p-tau as a biomarker associated with non-Alzheimer age-related cognitive decline. Several studies have reported significant negative associations between plasma p-tau181 and cross-sectional cognition [17–23]. The use of the MMSE in many of these studies may however lead to an underestimation of the relationship between cognitive function and plasma p-tau181 as the MMSE exhibits a ceiling effect in those without dementia [24] and exhibits restricted ageing trends [25] at odds with reports using psychometric psychological measurement [26]. Associations between p-tau181 and longitudinal cognitive ageing differences require more detailed psychometric measurement to characterise differences more accurately among non-demented older adults. Further, longitudinal data is important to fully reflect the within-person dynamics of the cognitive ageing phenomenon.

One longitudinal study using data from ADNI demonstrated a negative association between baseline plasma p-tau181 and 5-year cognitive decline in a memory composite in those cognitively unimpaired and those with MCI [27]. Clark and colleagues similarly found an association between baseline plasma p-tu181 and MMSE change over ~ 33 months change in those with cognitive impairment, but not those cognitively unimpaired in another cohort [28]. Finally, baseline plasma p-tau181 was found to be correlated with 1-year MMSE decline in a pooled sample of AD, MCI, and cognitively unimpaired participants, although this relationship was no longer significant when analysed by group separately. Despite the growing literature of longitudinal associations between plasma p-tau181 and cognitive function, few studies use participant-appropriate cognitive testing which is required to identify early and potentially subtle differences in cognitive functioning. This may account for most extant studies finding no association in those without cognitive impairment. In the current study, we examine this association in the LBC1936. This cohort offers the benefit of a narrow age range with a long follow-up period (~ 10 years) which provides a focused insight into those who are exclusively in the transition phase from 70 years old to 80 years, when the risk of cognitive decline starts to accelerate markedly [29].

Although research around plasma p-tau181 as a biomarker associated with cognitive function is burgeoning, relatively few studies have examined this phospho-site in brain tissue. Synapse loss is the strongest pathological correlate of cognitive decline in AD and region-specific synapse changes are also thought to contribute to age-related cognitive decline [30]. Emerging evidence suggests that abnormal tau accumulation in synapses contributes to synapse degeneration [31]. Previous work has reported evidence of p-tau accumulation in both AD and age-matched control brain tissue, specifically p-tau S202/Thr205 in crude homogenates [15] and S396/404 and S202 in synaptoneurosomes fractions [32]. It is currently unclear, however, whether p-tau181 is present in brain tissue, whether it is present in synapses, and whether there is a relationship between brain/synapse levels and plasma levels of the protein.

In the current study, we examined whether plasma levels of p-tau181 are associated with cognitive decline in a large cohort of older adults without dementia, the LBC1936. We hypothesised that greater plasma p-tau181 will be significantly associated with cognitive decline as measured by longitudinal changes in participant-suitable measure of cognitive function, the general cognitive factor (usually known as the *g* factor). A second aim of the current study is to examine whether p-tau181 is present in synapses in postmortem brain tissue and whether this is associated with cognitive decline.

## 2. Methods

### 2.1. Participants

#### 2.1.1. Plasma p-tau LBC1936 sample

Plasma p-tau181 data and *APOE* genotypes were obtained from 200 participants in a longitudinal study of ageing, the Lothian Birth Cohort 1936 study (LBC1936[33–35]). P-tau data in the current study were obtained at age 72 (*M* age = 72.46, *SD* = 0.70) and age 82 (*M* age = 82.03, *SD* = 0.46). Inclusion criteria included having complete data available at both ages for cognition. Participants were not eligible for selection if there was evidence of neuroradiologically-identified stroke. Of the 200 participants selected, five participants were diagnosed with dementia by wave 5 follow-up and were removed from analyses. Ethical approval was obtained from Multi-Centre Research Ethics Committee for Scotland (MREC/01/0/56; Wave 1), the Lothian Research Ethics Committee (LREC/2003/2/29; Wave 1), and the Scotland A Research Ethics Committee (07/MRE00/58; Waves 2–5).

#### 2.1.2. Post-mortem samples

Use of human tissue for post-mortem studies has been reviewed and approved by the Academic and Clinical Central Office for Research and Development medical research ethics committee (approval 15-HV-016) and the Edinburgh Brain Bank (research ethics committee approval 16/ES/0084). Tissue from 33 donors was used in the current study (Table 1). Donors in the western blot sample were either healthy ageing participants in the LBC1936 study (*n* = 12; referred to as healthy agers (HA)), people who died in mid-life with no known neurological or psychiatric disorders (*n* = 10; mid-life, ML), or people who died with both clinical and neuropathological diagnosis of Alzheimer’s disease (*n* = 11; AD). Data were available for donors in the HA group relating to pre-morbid lifetime cognitive decline computed from the MHT at age 11 and at ages 70-76. MHT at age 70-76 was regressed onto MHT at age 11; participants with positive residuals were classified as lifetime cognitive resilient (*n* = 5; LCR) as their MHT at age 70-76 is above what would be expected based on age 11 MHT. Those with negative residuals were classified as experiencing lifetime cognitive decline (*n* = 6; LCD). Twenty-one donors were included in the array tomography sample and were either HA (*n* = 16) or AD (*n* = 5).

**Table 1:** Post-mortem sample subject characteristics

| BBN number | Group | Method | Age | PMI | Sex | Braak stage | Thal Stage | Cognitive status |
| --- | --- | --- | --- | --- | --- | --- | --- | --- |
| 001.28406 | HA | WB<br>AT | 79 | 72 | Male | 2 | 2 | LCD |
| 001.32577 | HA | WB<br>AT | 81 | 74 | Male | 2 | 3 | LCD |
| 001.28793 | HA | WB<br>AT | 79 | 72 | Female | 2 | 1 | N/A |
| 001.28794 | HA | WB<br>AT | 79 | 61 | Female | 1 | 0 | LCR |
| 001.26495 | HA | WB<br>AT | 78 | 39 | Male | 1 | 1 | LCD |
| 001.28797 | HA | WB<br>AT | 79 | 57 | Male | 0 | 0 | LCR |
| 001.31495 | HA | WB<br>AT | 81 | 38 | Male | 6 | 4 | LCR |
| 001.28402 | HA | WB<br>AT | 79 | 49 | Male | 1 | 2 | LCD |
| 001.19686 | HA | WB<br>AT | 77 | 75 | Female | 1 | 1 | LCD |
| 001.34131 | HA | WB<br>AT | 82 | 95 | Male | 4 | 3 | LCD |
| 001.29082 | HA | WB<br>AT | 79 | 80 | Female | 3 | 5 | LCR |
| 001.29086 | HA | WB<br>AT | 79 | 68 | Female | 0 | 1 | LCR |
| 001.35215 | HA | AT | 82 | 40 | Male | 1 | 0 | LCR |
| 001.35549 | HA | AT | 82 | 56 | Male | 1 | 1 | LCR |
| 001.35866 | HA | AT | 83 | 78 | Female | 4 | 4 | LCD |
| 001.36135 | HA | AT | 84 | 30 | Female | 2 | 1 | LCR |
| 001.24479 | ML | WB | 46 | 76 | Female | 0 | 0 | N/A |
| 001.29906 | ML | WB | 51 | 52 | Male | N/A | N/A | N/A |
| 001.33613 | ML | WB | 46 | 99 | Female | 0 | 0 | N/A |
| 001.30169 | ML | WB | 48 | 58 | Male | N/A | N/A | N/A |
| 001.34244 | ML | WB | 49 | 69 | Female | 0 | 0 | N/A |
| 001.29466 | ML | WB | 39 | 76 | Male | 0 | 0 | N/A |
| 001.24342 | ML | WB | 33 | 47 | Male | 0 | 0 | N/A |
| 001.28792 | ML | WB | 58 | 51 | Male | N/A | N/A | N/A |
| 001.30972 | ML | WB | 34 | 99 | Male | 0 | 0 | N/A |
| 001.26976 | ML | WB | 19 | 101 | Male | 0 | 0 | N/A |
| 001.36066 | AD | WB | 94 | 29 | Male | 6 | 5 | N/A |
| 001.29695 | AD | WB<br>AT | 86 | 72 | Male | 6 | 5 | N/A |
| 001.35183 | AD | WB | 74 | 75 | Male | 6 | 5 | N/A |
| 001.30142 | AD | WB | 88 | 112 | Female | 6 | 5 | N/A |
| 001.29135 | AD | WB | 90 | 73 | Male | 6 | 3 | N/A |
| 001.35564 | AD | WB | 90 | 52 | Female | 6 | 5 | N/A |
| 001.30883 | AD | WB | 61 | 69 | Female | 6 | 5 | N/A |
| 001.29521 | AD | WB | 95 | 96 | Male | 6 | 5 | N/A |
| 001.30973 | AD | WB | 89 | 96 | Female | 6 | 5 | N/A |
| 001.36328 | AD | WB | 71 | 81 | Male | 6 | 5 | N/A |
| 001.36346 | AD | WB | 90 | 60 | Female | 6 | 5 | N/A |
| 001.32929 | AD | AT | 85 | 80 | Female | 6 | 5 | N/A |
| 001.35535 | AD | AT | 83 | 95 | Female | 6 | 4 | N/A |
| 001.35811 | AD | AT | 83 | 72 | Male | 6 | 4 | N/A |
| 001.26718 | AD | AT | 78 | 74 | Male | 6 | 5 | N/A |

Abbreviations: BBN, brain bank number; AD, Alzheimer's disease; AT, array tomography;
WB, western blot; *APOE*, apolipoprotein E; HA, healthy ager; ML, mid-life; LCD, lifetime cognitive decline; LCR, lifetime cognitive reserve; PMI, post-mortem interval (hours).

### 2.2. Immunoblotting

Brain homogenates and synaptoneurosomes were prepared according to [32]. 200mg of freshly frozen human brain tissue (BA20/21) was homogenised in 1mL buffer (25mM/L HEPES pH 7.5, 120mM/L NaC1, 5mM/L KCL 1mM/L MgC1_2_, 2mM/L CaC1_2_), with protease inhibitors (Roche complete mini) and phosphatase inhibitors (Millipore, Watford, UK). The homogenate was passed through an 80μm nylon filter (Millipore, Watford, UK) and a 300μL aliquot was saved and mixed with buffer (100mM/L Tris-HC1 pH 7.6, 4% SDS, protease inhibitor cocktail EDTA-free 100x Thermo Fisher Scientific, Loughborough, UK) to prepare the crude homogenate. The remainder of the homogenate was passed through a 5μm filter (Millipore, Watford, UK) then centrifuged at 1000 x g for 5 minutes. The supernatant was discarded and the pellet was washed with buffer and centrifuged again, yielding the synaptoneurosome pellet. Protein concentrations were determined using a protein assay (Thermo Fisher Scientific, Loughborough, UK). 20μg of protein per sample was electrophoresed in 4-12% Bis-Tris polyacrylamide gels (Invitrogen, Paisley, UK). Proteins were electro-transferred to nitrocellulose membranes (Thermo Fisher Scientific, Loughborough, UK) using the iBlot™ Dry Blotting system (#IB21001, Invitrogen, Paisley, UK). Revert 700 Total Protein Stain was used to quantify total protein (Li-Cor, Cambridge, UK). Membranes were incubated in block buffer, then incubated in primary antibodies in blocking buffer (p-tau181 (1:500, Invitrogen AT270 #MN1050) and total tau (1:500, Tau13 BioLegend #MMS-520R)). Membranes were washed and incubated with secondary antibodies (1:5000, Li-Cor Biosciences), rinsed and imaged using the Odyssey Imaging system, and analysed using Open Image Studio Lite.

### 2.3. Array Tomography

Samples from dorsolateral prefrontal cortex were trimmed into blocks and fixed in 4% paraformaldehyde for 3 hours [36]. Samples were dehydrated, embedded in LR White resin, and cut into ribbons of 70nm serial sections. Ribbons were rehydrated with 50mM glycine in TBS, washed in TBS, and incubated in blocking buffer (0.05% Tween, 0.1% fish skin gelatine in TBS) followed by primary antibodies (p-tau181 (1:100, AT270), PSD-95 (1:50, Cell Signalling #3450), and Synaptophysin (1:100, R&D Systems #AF5555). Secondary antibodies were diluted in block buffer (1:50) and incubated for 45 minutes at room temperature (goat anti-guinea pig IgG H&L Cy3 (Abcam #102370), and donkey anti-mouse IgG H&L Alexa Fluor 647 (Abcam #150107). Alexa Fluor 488 labelled anti-synaptophysin antibody (1:100, Abcam #196379) was then applied for 1 hour after initial secondary antibodies. Images were acquired on a Leica TCS SP8 confocal and analysed using FIJI, MATLAB (version 2018a) and Docker to run python-based analysis scripts, which are freely available on GitHub https://github.com/arraytomographyusers/Array_tomography_analysis_tool.

### 2.4. Computation of change in general cognitive function

Participants in the LBC1936 study were given a detailed battery of neuropsychological tests at waves 2 (age ~ 73), 3 (age ~ 76), 4 (age ~ 79), and 5 (age ~ 82). In the current study, we used the following six non-verbal subtests from the Wechsler Adult Intelligence Scale-III UK (WAIS-III) [37] to compute a slope representing general fluid cognitive (*g*) decline (from here referred to as *g* factor change): matrix reasoning, letter-number sequencing, block design, symbol search, digit symbol coding, and digit span backwards. Individual scores of cognitive ageing were computed for the total LBC1936 sample by fitting a Factor of Curves model in a structural equation modelling (SEM) framework using full information maximum likelihood estimation in R, using the *lavaan* package [38]. The latent slope scores were extracted for further analysis..

### 2.5. P-tau181 plasma assay

Plasma assays were conducted at the biomarker lab at the UK Dementia Research Institute at UCL. Plasma p-tau181 (*i.e.*, tau phosphorylated at threonine 181) concentration was measured using the Simoa® pTau-181 Advantage Kit on an HD-X Analyzer (Quanterix, Billerica, MA). Plasma samples were thawed and centrifuged at 13000 ×◻g for 5◻min at room temperature. Calibrators (neat) and samples (plasma: 1:4 dilution) were measured in duplicates. Assays were performed using the Simoa HD-X Analyzer (Quanterix Corp, Billerica, MA, USA). All samples were analysed using the same batch of reagents. A four-parameter logistic curve fit data reduction method was used to generate a calibration curve. Two control samples of known concentration of the protein of interest (high-ctrl and low-ctrl) were included as quality control. Mean coefficient of variation was 8.29%.

### 2.6. Statistical Analysis

R Studio (version 4.0.4) was used for all data analysis. Data were tested for normality by inspection of histograms. Group differences were analysed using *t*-tests, Wilcoxon rank-sum, and Kruskal-Wallis where appropriate. Correlations were conducted using Pearson’s r or Spearman’s rho dependent on normality of data. Cohen’s *d* was used as a measure of effect size [39]. Wilcoxon rank-sum tests were used to examine group differences in brain p-tau181. Healthy agers were split into those with age-related cognitive decline and those with maintained cognition and Wilcoxon rank-sum tests were used to examine group differences in TH and SN p-tau181.

To examine whether baseline plasma p-tau181 could predict change in cognitive function, two models were fitted. The first where *g* factor change was regressed onto sex, baseline age (years), and baseline p-tau181. The second model added education (years), age 11 IQ, and *APOE* status. Baseline p-tau181 values were log-transformed to adjust for a non-normal distribution. Change in serum p-tau181 was computed by regressing values at wave 5 onto values at wave 2. The standardised residuals from this model were then entered into identical models as above. Standardized coefficients are reported (β) with bootstrapped 95% confidence intervals (CI) from 1000 replications. Significance was reported when *P* < .05.

## 3. Results

### 3.1. Plasma p-tau181 predicts cognitive decline

In total 195 participants were included in analyses with baseline p-tau181, and 192 in analysis examining p-tau181 change. Cognitive and plasma-based data were available for two time-points when participants were a mean age of 72.46 (*SD* = 0.70) and 82.02 years (*SD* = 0.46). Demographic information is provided in Table 2. Descriptive plots of p-tau181 change are shown in Figure 1A–1C. P-tau181 levels were significantly elevated at wave 5 relative to wave 2 (*W* = 12867, *P* < .01, *d* = 0.30, 95% CI = 0.15 – 0.44). Males and females differed significantly in plasma p-tau181 levels at wave 2 (*W* = 5487, *P* = .03, *d* = 0.37, 95% CI = 0.09 – 0.65), but not at wave 5 (*W* = 4893, *P* = .44). Age at wave 2 and p-tau181 levels at wave 2 were significantly correlated across the cohort (*rho* = 0.23, *P* = <.01), however, the relationship between age at wave 5 and p-tau181 at wave 5 did not reach significance (*rho* = 0.13, *P* = .08). At wave 2, there were no significant differences in p-tau between *APOE* ε4 carriers and non-carriers (*W* = 2868, *P* = .11), however, at wave 5 *APOE* ε4 carriers had significantly elevated levels (*W* = 2518, *P* < .01; *d* = 0.25, 95% CI = 0.07 – 0.57; see Figure 1D).

**Table 2:** Plasma p-tau LBC1936 subject characteristics

| Variable | Mean (SD) |
| --- | --- |
| Wave 2 Age (Years) | 72.46 (0.70) |
| Wave 5 Age (Years) | 82.02 (0.46) |
| Wave 2 Plasma p-tau181 pg/mL | 24.74 (13.35) |
| Wave 5 Plasma p-tau181 pg/mL | 30.34 (18.62) |
| Age 11 IQ | 103.83 (14.04) |
| Education (Years) | 11.01 (1.16) |
|  | *n* (%) |
| Sex (F:M) | 102:93 |
| *APOE* ε4 carriers | 52(26.67%) |

Abbreviations: p-tau181, phosphorylated tau 181; IQ, intelligence quotient; *APOE*, apolipoprotein E

**Figure 1:**
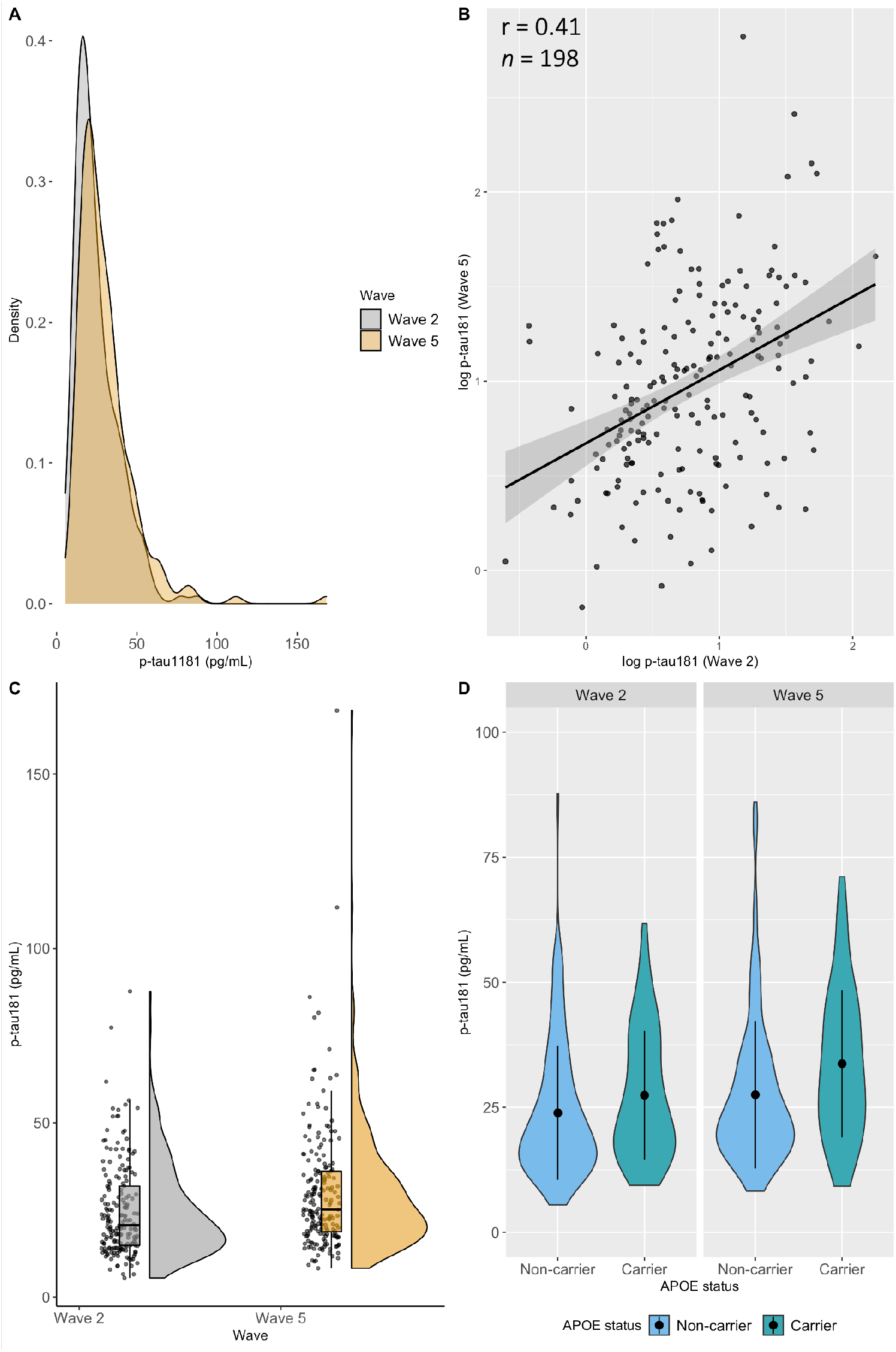
(A) Density plot of plasma p-tau181 at wave 2 and wave 5; (B) scatterplot of log-transformed plasma p-tau181 at wave 2 and wave 5 with regression line and 95% confidence intervals. Pearson correlation denoted by r, sample size denoted by *n*; (C) Raincloud plot[49,50] to show distribution of p-tau181 across wave 2 and wave 5; (D) Violin plot showing mean and distribution of plasma p-tau181 at wave 2 and wave 5 between *APOE* ε4 carriers and non-carriers.

In model 1 (baseline p-tau181, age, sex), p-tau181 significantly predicted *g* factor decline (β = −0.17, 95% CI = −0.15 – −0.01, *P* = .02), where a 1 SD higher baseline level of p-tau181 is associated with 0.17 SD decrease in *g* factor. In model 2 (addition of age 11 IQ, education years, *APOE* status), baseline p-tau181 significantly predicted *g* factor decline (β = −0.17, 95% CI = −0.16 – −0.01, *P* = .03). Next, *g* factor change was regressed onto p-tau181 change, sex, and age. P-tau181 change was a significant predictor of *g* factor decline (β = −0.23, 95% CI = −0.09 – −0.02, *P* = <.01). When including covariates age 11 IQ, education years, and *APOE* status, p-tau181 change remained a significant predictor of *g* factor decline (β = −0.21, 95% CI = −0.08 – −0.01, *P* = <.01), where a 1 SD increase in p-tau181 change was associated with a 0.21 SD decrease in *g* factor.

### 3.2. Post-mortem analyses show p-tau181 in synapses

Thirty-three cases were used for western blot analyses of brain tissue, with 12 HA, 10 mid-life, and 11 AD cases. Individual case details are provided in Table 1, sample summary statistics are provided in Table 3. Between the three groups, there were no significant differences in sex (c^2^, *df* = 2, *P* = .75) or in PMI (*H* = 0.61, *df* = 2, *P* = .73). Both AD and HA donors were significantly older than mid-life donors (*H* = 41.91, *df* = 2, *P < .01)*, although there was no significant difference between AD and HA donors (*P* = .06).

**Table 3:** Post-mortem sample summary statistics

| Variable | HA (*n* = 12) | ML (*n* = 10) | AD (*n* = 11) |
| --- | --- | --- | --- |
| Sex (F:M) | 5:7 | 3:7 | 5:6 |
| Age (years) | 79.3 (1.34)<sup>†</sup> | 42.30 (10.94)<sup>‡</sup> | 84.36 (10.56) |
| PMI (hours) | 65 (16.56) | 72.80 (20.51) | 74.09 (22.25) |

Abbreviations: AD, Alzheimer's disease; HA, healthy ager; ML, mid-life; PMI, post-mortem interval. \* indicates significant difference compared to both groups, <sup>†</sup> versus ML group, <sup>‡</sup> versus AD group

As seen in figure 2A, across the three groups, there were no significant differences in total tau levels (*H* = 6.12, *df* = 2, *P* = .05) in total homogenate, nor in synaptoneurosome preparations (*H* = 0.83, *df* = 2, *P* = .66; figure 2B). When the HA sample was split by premorbid cognitive status, there were no significant differences in total tau in either total homogenate (*W* = 20, *P* = .11) or synaptoneurosome preparations (*W* = 9, *P* = .99).

**Figure 2:**
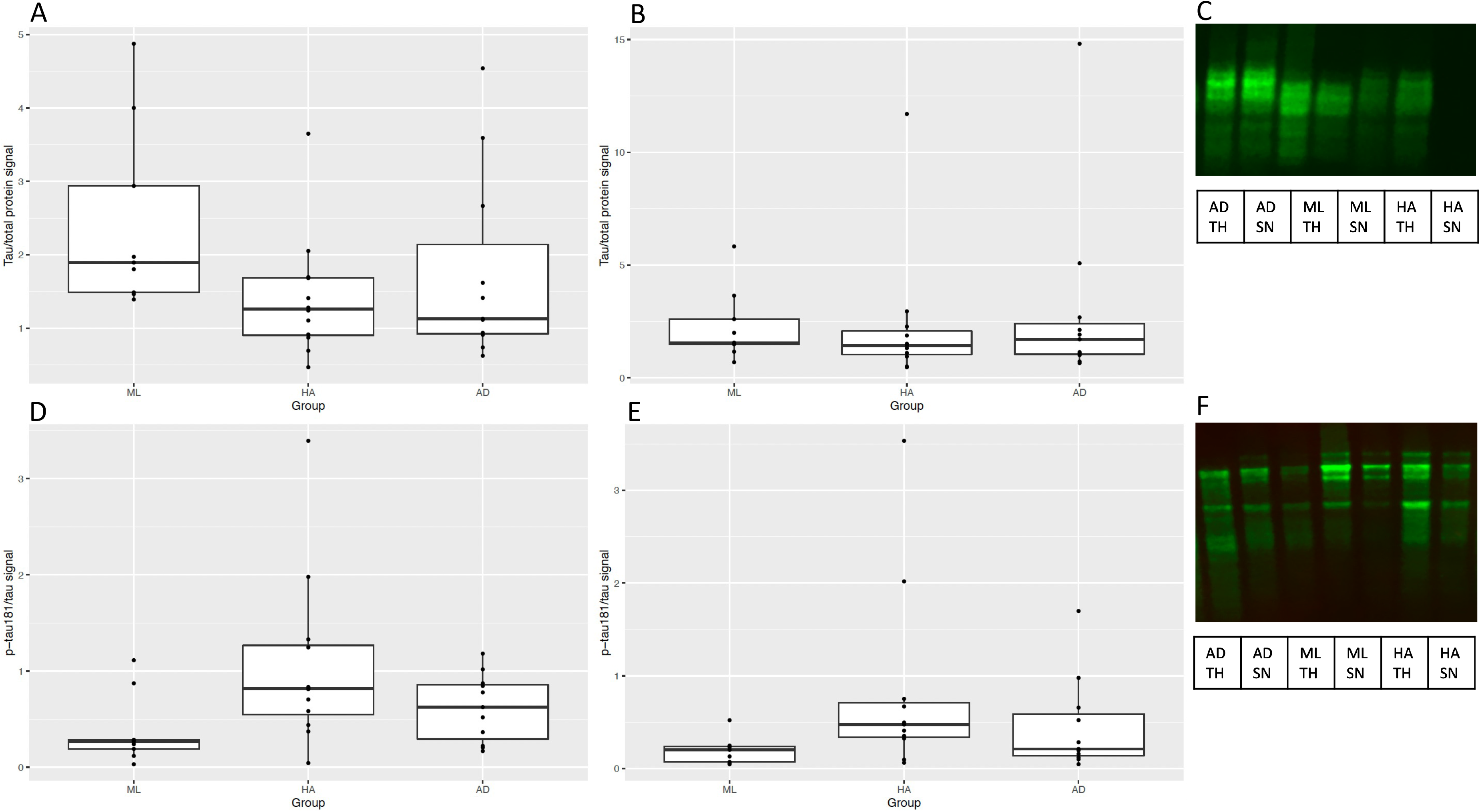
Boxplots of total tau in total homogenate (A) and synaptoneurosomes (B). Western blot show in in (C). Boxplots of p-tau181 in total homogenate (D) and synaptoneurosomes (E). Western blot shown in (F). Total tau and p-tau181 were stained on separate gels. AD = Alzheimer’s disease; ML = mid-life; HA = healthy agers; TH = total homogenate; SN = synaptoneurosome.

In total homogenate preparations, there were no significant differences in p-tau181 across the three groups (Figure 2D; *H* = 5.59, *df* = 2, *P* = .06), nor in synaptoneurosome preparations (*H* = 5.38, *df* = 2, *P* = .07; Figure 2E). There were no significant differences in p-tau181 between the HA sub-groups of lifetime cognitive resilience or lifetime cognitive decline in either total homogenate (*W* = 6, *P* = .26) or synaptoneurosome preparations (*W* = 10, *P* = .90).

Synaptoneurosome preparations contain both pre and post-synaptic elements and cannot distinguish which side of the synapse contains ptau-181. High resolution array tomography imaging was used to examine whether ptau-181 is detectable in pre and/or post-synapses in human brain tissue. In blocks of dorsolateral prefrontal cortex samples from 10 LBC1936 participants and five people with AD, we observe abundant ptau-181 staining in processes and localisation in both pre and post synapses (Figure 3). Consistent with the western blot synaptoneurosome data, there are no qualitative differences between LBC1936 participants who exhibited cognitive decline compared to those who had maintained cognition.

**Figure 3:**
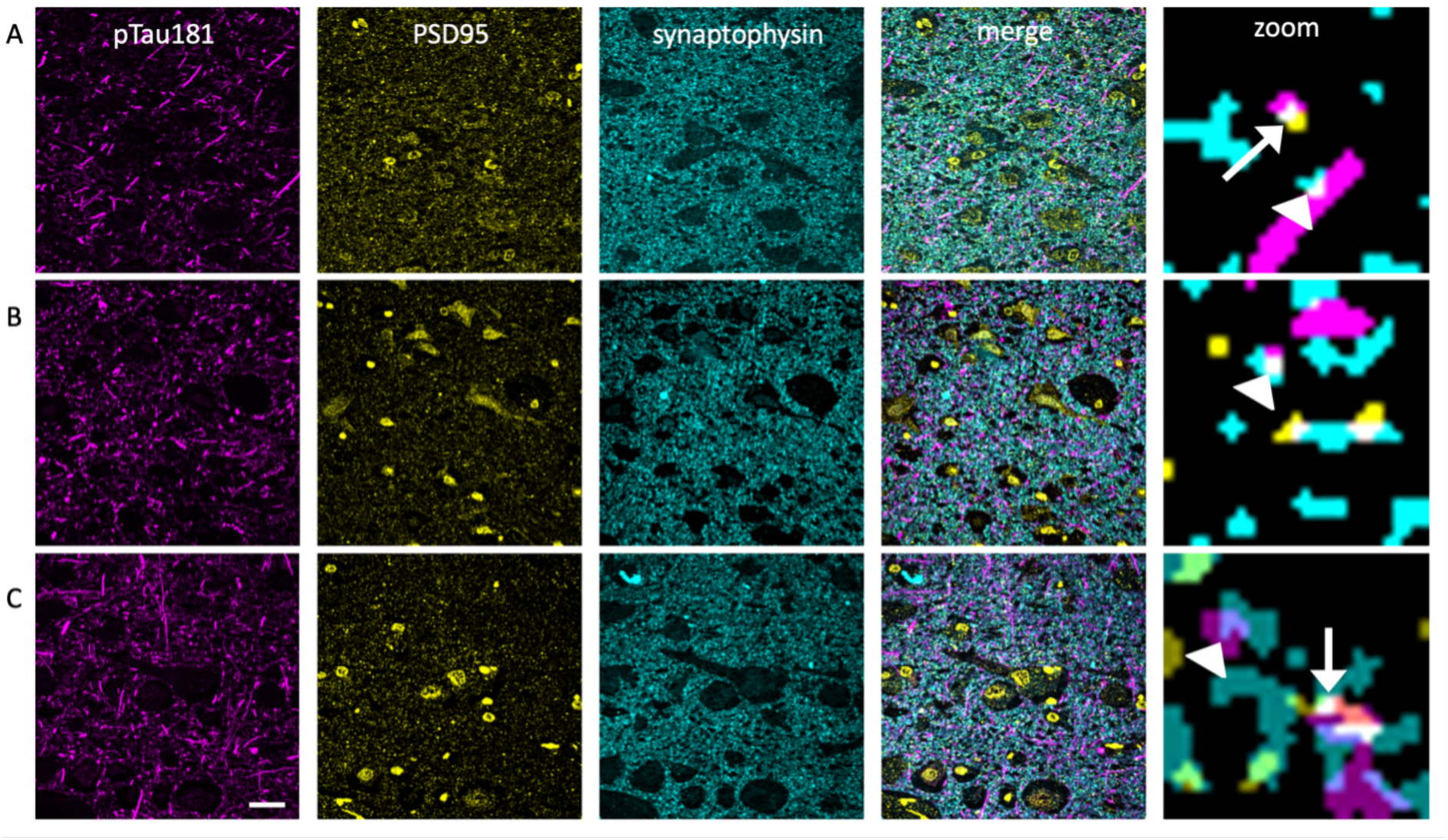
Array tomography imaging of a LBC1936 participant with maintained cognition (A), LBC1936 participant with age-related cognitive decline (B) and an Alzheimer’s disease subject (C) shows ptau-181 (magenta) in neurites and present in a subset of post-(yellow) and pre-(cyan) synapses. Images in the first three columns show maximum intensity z-projections of 10 serial sections (scale bar 20 mm). The far-right column shows a 10 mm x 10 mm zoom of a single 70 nm section after segmentation to detect synapses. In the zoomed images, ptau-181 is colocalised with presynaptic terminals (arrowheads) and post-synaptic terminals (arrows).

## 4. Discussion

In the current study, we identified a significant association between cognitive ageing and both baseline plasma p-tau181 and p-tau181 change. That is, individuals with elevated baseline p-tau181 had greater cognitive decline ~10 years later. Further, those a greater increase in plasma p-tau181 over ~10 years had greater cognitive decline over the same period. We also found that with increasing age, *APOE* ε4 carriers showed significantly higher p-tau18. Associations between p-tau181 and *g* factor decline were corrected for *APOE* status, one of the predictors of cognitive ageing in this and other samples [40–43], and remained significant. This highlights the unique potential value of this marker in accounting for differences in cognitive ageing among community-dwelling older adults. In brain p-tau181, we found no significant differences in either total homogenate or synaptoneurosome preparations across HA, ML, and AD cases. Finally, in the current study we also report evidence of the presence of p-tau181 in synapses.

Our findings suggest that plasma p-tau181 may be informative about cognitive decline in those without a diagnosis of dementia. Previous work has also reported significant associations between plasma p-tau181 and cognitive decline across the AD-spectrum, however, much of the extant literature around those with no cognitive impairment reported no association with longitudinal measures. Our findings contrast with much of the existing literature, however, in the current study we used a more sensitive and participant-appropriate measure of cognitive functioning associated with ageing. Furthermore, our sample has the benefit of a narrow age range which offers a more precise insight into those transitioning the 8^th^ to 9^th^ decades of life.

Our study is one of the few studies examining plasma p-tau181 and cognitive function longitudinally, and one of the first to do so in an aged cohort without dementia over a ~ 10-year period. Our advanced statistical treatment of cognitive change, which was based on measurements at four occasions allowed us to reliably estimate cognitive change, and more precisely partition the variance attributable to cognitive declines as distinct from extant differences in prior ability. Moreover, this is the first known study to examine the relationship between change in plasma p-tau181 and change in cognitive function. Taken together, this suggests that p-tau181 is present during typical ageing and so may be a suitable marker for age-related cognitive decline.

We observe ptau-181 within synapses in human brain, where it may contribute to cognitive function or cognitive decline. However, we found no differences in ptau-181 in total brain homogenate or synaptoneurosome preparations between AD, mid-life, and healthy agers. To our knowledge, this is the first study to examine p-tau181 in synapses, however, previous work has reported elevations of p-tau S396/404 and S202 in AD synaptoneurosomes [32]. The finding of no differences in our study may be partially explained by a lack of power. Our sample size was relatively small which may increase the chances of a type II error. Where post-mortem data are a precious resource, especially with pre-morbid cognitive function data, we acknowledge that a larger sample size would enable more reliable detection and estimation of effect sizes.

The association between plasma p-tau181 and cognitive decline in typical ageing suggests it may be a suitable marker for age-related cognitive decline. This could become an economical and time-efficient screening tool for the evaluation of individuals who are most at-risk of cognitive decline, and thus are most eligible for clinical trials. Our results suggest that baseline p-tau181 and p-tau181 have similar magnitudes of effect of cognitive decline. Looking at the standardised coefficients in the current study, there is an increase in the steepness of 0.17 SDs in cognitive function decline for every 1 SD increase of baseline p-tau181. Further, there is an increase in the steepness of 0.21 SDs in cognitive function decline for every 1 SD increase in p-tau181 over time. Our measure of change was statistically independent of baseline p-tau181, suggesting both measures are important for predicting cognitive decline. That is, those with higher baseline p-tau181 and a steeper increase over time (irrespective of baseline), are also those with steeper cognitive decline.

This study is not without limitations. Participants were selected for analyses if complete cognitive test and plasma samples were available at ages 72 and 82 so as to maximise the power of our longitudinal analyses. This could lead to a selection bias impacting results as we reported significant differences in education, age 11 IQ, and *g* factor change between those included and those not included. However, prior work in this cohort indicates that the studies’ initial selection bias and subsequent pattern of attrition skews our design toward healthier and more-educated individuals, and that our results therefore likely underestimate effects present at the population level [35,44]. Furthermore, it is unknown when plasma p-tau181 increases begin during the lifetime. The current sample is comprised of individuals in later-life. Future longitudinal research examining p-tau181 from mid-life onwards could be useful in understanding when plasma p-tau181 increases begin and whether cognitive decline can be predicted at an earlier age. Finally, LBC1936 participants with plasma p-tau181 data were excluded if they had a diagnosis of dementia, however, this does not guarantee that the sample is “typically ageing” and the presence of early neurodegenerative processes cannot be ruled out.

It will also be important to investigate differences which may affect the clinical implementation of plasma biomarkers, such as sex and ethnicity differences, as well as lifestyle factors associated with p-tau levels. For example, cognitive impairment is associated with smaller changes in CSF tau in an African-American sample relative to a Caucasian sample [45]. Furthermore, more studies are needed to compare plasma p-tau181 as a biomarker with more established biomarkers such as tau PET, MRI measures, and CSF tau. Quantification of p-tau181 at the level of the synapse using AT, as well as larger samples for immunoblotting would benefit the field in characterising p-tau181 in the brain and future work would benefit from examining correlations between brain p-tau181 and plasma p-tau181, which was not possible in the current study. Finally, the investigation of p-tau phosphorylated at other sites should also be investigated; plasma p-tau231 and p-tau217 have both been investigated as promising biomarkers associated with cognitive function [9,46–48]

The current study provides the first evidence in an exclusively older-age sample with narrow age-range that both the level and subsequent change in plasma p-tau181 are associated with subsequent cognitive decline between 73-82 years, beyond *APOE* status and in the absence of dementia. We also report evidence that it is present in synapses where it may influence cognition. Overall, these findings suggest p-tau181 may be associated with age-related cognitive decline.

## References

[1] World Health Organisation. Ageing and health factsheet. 2018 n.d. https://www.who.int/news-room/fact-sheets/detail/ageing-and-health (accessed August 25, 2021).

[2] Tucker-Drob EM, Briley DA, Starr JM, Deary IJ. Structure and Correlates of Cognitive Aging in a Narrow Age Cohort. Psychol Aging 2014;29:236. https://doi.org/10.1037/A0036187.

[3] Bárrios H, Narciso S, Guerreiro M, Maroco J, Logsdon R, de Mendonça A. Quality of life in patients with mild cognitive impairment. Aging Ment Health 2013;17:287–92. https://doi.org/10.1080/13607863.2012.747083.

[4] Boyle P, Wilson R, Yu L, Barr A, Honer W, Schneider J, et al. Much of late life cognitive decline is not due to common neurodegenerative pathologies. Ann Neurol 2013;74:478–89. https://doi.org/10.1002/ANA.23964.

[5] Arendt T, Stieler J, Holzer M. Tau and tauopathies. Brain Res Bull 2016;126:238–92. https://doi.org/10.1016/J.BRAINRESBULL.2016.08.018.

[6] Ballatore C, Lee V, Trojanowski J. Tau-mediated neurodegeneration in Alzheimer’s disease and related disorders. Nat Rev Neurosci 2007;8:663–72. https://doi.org/10.1038/NRN2194.

[7] Zetterberg H, Burnham SC. Blood-based molecular biomarkers for Alzheimer’s disease. Mol Brain 2019;12. https://doi.org/10.1186/s13041-019-0448-1.

[8] Dhiman K, Blennow K, Zetterberg H, Martins RN, Gupta VB. Cerebrospinal fluid biomarkers for understanding multiple aspects of Alzheimer’s disease pathogenesis. Cell Mol Life Sci 2019;76:1833–63. https://doi.org/10.1007/s00018-019-03040-5.

[9] Thijssen EH, Joie R La, Strom A, Fonseca C, Iaccarino L, Wolf A, et al. Plasma phosphorylated tau 217 and phosphorylated tau 181 as biomarkers in Alzheimer’s disease and frontotemporal lobar degeneration: a retrospective diagnostic performance study. Lancet Neurol 2021;20:739–52. https://doi.org/10.1016/S1474-4422(21)00214-3.

[10] Thijssen E, La Joie R, Wolf A, Strom A, Wang P, Iaccarino L, et al. Diagnostic value of plasma phosphorylated tau181 in Alzheimer’s disease and frontotemporal lobar degeneration. Nat Med 2020;26:387–97. https://doi.org/10.1038/S41591-020-0762-2.

[11] S J, N M, S P, R S, Tg B, Ge S, et al. Plasma P-tau181 in Alzheimer’s disease: relationship to other biomarkers, differential diagnosis, neuropathology and longitudinal progression to Alzheimer’s dementia. Nat Med 2020;26:379–86. https://doi.org/10.1038/S41591-020-0755-1.

[12] Lantero Rodriguez J, Karikari TK, Suárez-Calvet M, Troakes C, King A, Emersic A, et al. Plasma p-tau181 accurately predicts Alzheimer’s disease pathology at least 8 years prior to post-mortem and improves the clinical characterisation of cognitive decline. Acta Neuropathol 2020;140:267–78. https://doi.org/10.1007/s00401-020-02195-x.

[13] Chen T Bin, Lai YH, Ke TL, Chen JP, Lee YJ, Lin SY, et al. Changes in Plasma Amyloid and Tau in a Longitudinal Study of Normal Aging, Mild Cognitive Impairment, and Alzheimer’s Disease. Dement Geriatr Cogn Disord 2020;48:180–95. https://doi.org/10.1159/000505435.

[14] Negash S, Bennett DA, Wilson RS, Schneider JA, Arnold SE. Cognition and Neuropathology in Aging: Multidimensional Perspectives from the Rush Religious Orders Study and Rush Memory and Aging Project. Curr Alzheimer Res 2011;8:336.

[15] Tzioras M, Easter J, Harris S, McKenzie C-A, Smith C, Deary I, et al. Assessing amyloid-β, tau, and glial features in Lothian Birth Cohort 1936 participants post-mortem. Matters 2017;3:e201708000003. https://doi.org/10.19185/matters.201708000003.

[16] Bennett DA, Wilson RS, Boyle PA, Buchman AS, Schneider JA. Relation of Neuropathology to Cognition in Persons Without Cognitive Impairment. Ann Neurol 2012;72:599. https://doi.org/10.1002/ANA.23654.

[17] Chen S-D, Huang Y-Y, Shen X-N, Guo Y, Tan L, Dong Q, et al. Longitudinal plasma phosphorylated tau 181 tracks disease progression in Alzheimer’s disease. Transl Psychiatry 2021;11. https://doi.org/10.1038/S41398-021-01476-7.

[18] Wang Y-L, Chen J, Du Z-L, Weng H, Zhang Y, Li R, et al. Plasma p-tau181 Level Predicts Neurodegeneration and Progression to Alzheimer’s Dementia: A Longitudinal Study. Front Neurol 2021;12:695696. https://doi.org/10.3389/FNEUR.2021.695696.

[19] Lussier FZ, Benedet AL, Therriault J, Pascoal TA, Tissot C, Chamoun M, et al. Plasma levels of phosphorylated tau 181 are associated with cerebral metabolic dysfunction in cognitively impaired and amyloid-positive individuals. Brain Commun 2021;3. https://doi.org/10.1093/BRAINCOMMS/FCAB073.

[20] Chong JR, Ashton NJ, Karikari TK, Tanaka T, Saridin FN, Reilhac A, et al. Plasma P-tau181 to Aβ42 ratio is associated with brain amyloid burden and hippocampal atrophy in an Asian cohort of Alzheimer’s disease patients with concomitant cerebrovascular disease. Alzheimer’s Dement 2021. https://doi.org/10.1002/ALZ.12332.

[21] Xiao Z, Wu X, Wu W, Yi J, Liang X, Ding S, et al. Plasma biomarker profiles and the correlation with cognitive function across the clinical spectrum of Alzheimer’s disease. Alzheimers Res Ther 2021;13:123. https://doi.org/10.1186/S13195-021-00864-X.

[22] Tsai C-L, Liang C-S, Lee J-T, Su M-W, Lin C-C, Chu H-T, et al. Associations between Plasma Biomarkers and Cognition in Patients with Alzheimer’s Disease and Amnestic Mild Cognitive Impairment: A Cross-Sectional and Longitudinal Study. J Clin Med 2019;8. https://doi.org/10.3390/JCM8111893.

[23] Karikari T, Pascoal T, Ashton N, Janelidze S, Benedet A, Rodriguez J, et al. Blood phosphorylated tau 181 as a biomarker for Alzheimer’s disease: a diagnostic performance and prediction modelling study using data from four prospective cohorts. Lancet Neurol 2020;19:422–33. https://doi.org/10.1016/S1474-4422(20)30071-5.

[24] Spencer R, Wendell C, Giggey P, Katzel L, Lefkowitz D, Siegel E, et al. Psychometric limitations of the mini-mental state examination among nondemented older adults: an evaluation of neurocognitive and magnetic resonance imaging correlates. Exp Aging Res 2013;39:382–97. https://doi.org/10.1080/0361073X.2013.808109.

[25] Matthews F, Marioni R, Brayne C. Examining the influence of gender, education, social class and birth cohort on MMSE tracking over time: A population-based prospective cohort study. BMC Geriatr 2012;12:1–5. https://doi.org/10.1186/1471-2318-12-45/FIGURES/2.

[26] Tucker-Drob EM. Cognitive Aging and Dementia: A Life-Span Perspective. Https://DoiOrg/101146/Annurev-Devpsych-121318-085204 2019;1:177–96. https://doi.org/10.1146/ANNUREV-DEVPSYCH-121318-085204.

[27] Therriault J, Benedet AL, Pascoal TA, Lussier FZ, Tissot C, Karikari TK, et al. Association of plasma P-tau181 with memory decline in non-demented adults. Brain Commun 2021;3. https://doi.org/10.1093/BRAINCOMMS/FCAB136.

[28] Clark C, Lewczuk P, Kornhuber J, Richiardi J, Maréchal B, Karikari TK, et al. Plasma neurofilament light and phosphorylated tau 181 as biomarkers of Alzheimer’s disease pathology and clinical disease progression. Alzheimers Res Ther 2021;13. https://doi.org/10.1186/S13195-021-00805-8.

[29] Plassman B, Lang K, Fisher G, Heeringa S, Weir D, Ofstedal M, et al. Prevalence of cognitive impairment without dementia in the United States. Ann Intern Med 2008;148:427–34. https://doi.org/10.7326/0003-4819-148-6-200803180-00005.

[30] Henstridge CM, Pickett E, Spires-Jones TL. Synaptic pathology: A shared mechanism in neurological disease. Ageing Res Rev 2016;28:72–84. https://doi.org/10.1016/J.ARR.2016.04.005.

[31] Spires-Jones TL, Hyman BT. The intersection of amyloid beta and tau at synapses in Alzheimer’s disease. Neuron 2014;82:756–71. https://doi.org/10.1016/j.neuron.2014.05.004.

[32] Tai H, Serrano-Pozo A, Hashimoto T, Frosch M, Spires-Jones T, Hyman B. The synaptic accumulation of hyperphosphorylated tau oligomers in Alzheimer disease is associated with dysfunction of the ubiquitin-proteasome system. Am J Pathol 2012;181:1426–35. https://doi.org/10.1016/J.AJPATH.2012.06.033.

[33] Deary IJ, Gow AJ, Taylor MD, Corley J, Brett C, Wilson V, et al. The Lothian Birth Cohort 1936: a study to examine influences on cognitive ageing from age 11 to age 70 and beyond. BMC Geriatr 2007;7:28. https://doi.org/10.1186/1471-2318-7-28.

[34] Deary IJ, Gow AJ, Pattie A, Starr JM. Cohort Profile: The Lothian Birth Cohorts of 1921 and 1936. Int J Epidemiol 2012;41:1576–84. https://doi.org/10.1093/IJE/DYR197.

[35] Taylor AM, Pattie A, Deary IJ. Cohort Profile Update: The Lothian Birth Cohorts of 1921 and 1936. Int J Epidemiol 2018;47:1042–1042r. https://doi.org/10.1093/ije/dyy022.

[36] Kay KR, Smith C, Wright AK, Serrano-pozo A, Pooler AM, Koffie R, et al. Studying synapses in human brain with array tomography and electron microscopy 2013;8:1366–80. https://doi.org/10.1038/nprot.2013.078.Studying.

[37] Wechsler D. Wechsler Memory Scale III-UK Administration and Scoring Manual. London: Psychological Corporation; 1998.

[38] Rosseel Y. lavaan: An R Package for Structural Equation Modeling. J Stat Softw 2012;48:1–36. https://doi.org/10.18637/JSS.V048.I02.

[39] Cohen J. Statistical Power Analysis for the Behavioral Sciences. New York, NY: Routledge Academic; 1988. https://doi.org/10.4324/9780203771587.

[40] Albrecht MA, Szoeke C, Maruff P, Savage G, Lautenschlager NT, Ellis KA, et al. Longitudinal cognitive decline in the AIBL cohort: The role of APOE ε4 status. Neuropsychologia 2015;75:411–9. https://doi.org/10.1016/J.NEUROPSYCHOLOGIA.2015.06.008.

[41] Davies G, Harris SE, Reynolds CA, Payton A, Knight HM, Liewald DC, et al. A genome-wide association study implicates the APOE locus in nonpathological cognitive ageing. Mol Psychiatry 2014 191 2012;19:76–87. https://doi.org/10.1038/mp.2012.159.

[42] Seeman TE, Huang MH, Bretsky P, Crimmins E, Launer L, Guralnik JM. Education and APOE-e4 in Longitudinal Cognitive Decline: MacArthur Studies of Successful Aging. Journals Gerontol Ser B 2005;60:P74–83. https://doi.org/10.1093/GERONB/60.2.P74.

[43] Schiepers OJG, Harris SE, Gow AJ, Pattie A, Brett CE, Starr JM, et al. APOE E4 status predicts age-related cognitive decline in the ninth decade: Longitudinal follow-up of the Lothian Birth Cohort 1921. Mol Psychiatry 2012;17:315–24. https://doi.org/10.1038/mp.2010.137.

[44] Johnson W, Brett CE, Calvin C, Deary IJ. Childhood characteristics and participation in Scottish Mental Survey 1947 6-Day Sample Follow-ups: Implications for participation in aging studies. Intelligence 2016;54:70–9. https://doi.org/10.1016/J.INTELL.2015.11.006.

[45] Howell JC, Watts KD, Parker MW, Wu J, Kollhoff A, Wingo TS, et al. Race modifies the relationship between cognition and Alzheimer’s disease cerebrospinal fluid biomarkers. Alzheimer’s Res Ther 2017;9. https://doi.org/10.1186/s13195-017-0315-1.

[46] Ashton N, Pascoal T, Karikari T, Benedet A, Lantero-Rodriguez J, Brinkmalm G, et al. Plasma p-tau231: a new biomarker for incipient Alzheimer’s disease pathology. Acta Neuropathol 2021;141:709–24. https://doi.org/10.1007/S00401-021-02275-6.

[47] Mattsson-Carlgren N, Janelidze S, Palmqvist S, Cullen N, Svenningsson AL, Strandberg O, et al. Longitudinal plasma p-tau217 is increased in early stages of Alzheimer’s disease. Brain 2020. https://doi.org/10.1093/brain/awaa286.

[48] Brickman AM, Manly JJ, Honig LS, Sanchez D, Reyes-Dumeyer D, Lantigua RA, et al. Plasma p-tau181, p-tau217, and other blood-based Alzheimer’s disease biomarkers in a multi-ethnic, community study. Alzheimer’s Dement 2021;17:1353–64. https://doi.org/10.1002/ALZ.12301.

[49] Allen M, Poggiali D, Whitaker K, Rhys Marshall T, Van Langen J, Kievit RA, et al. Raincloud plots: a multi-platform tool for robust data visualization. Wellcome Open Res 2021 463 2021;4:63. https://doi.org/10.12688/wellcomeopenres.15191.2.

[50] Whitaker K, TomRhys Marshall, Mourik T van, Martinez PA, poggiali davide, Ye H, et al. RainCloudPlots/RainCloudPlots: WellcomeOpenResearch 2019. https://doi.org/10.5281/ZENODO.3368186.

